# HIV-1 Protease Uses Bi-Specific S2/S2’ Subsites To Optimize Cleavage of Two Classes of Target Sites

**DOI:** 10.1101/404186

**Authors:** Marc Potempa, Sook-Kyung Lee, Nese KurtYilmaz, Ellen A. Nalivaika, Amy Rogers, Ean Spielvogel, Charles W. Carter, Celia A. Schiffer, Ronald Swanstrom

**Author notes:** Present address: Department of Microbiology and Immunology, University of California at San Francisco, San Francisco, CA, USA. Correspondence to Ronald Swanstrom: Department of Biochemistry and Biophysics, University of North Carolina at Chapel Hill, Chapel Hill, NC, 27599, USA.

## Abstract

Retroviral proteases (PR) have a unique specificity that allows cleavage of sites with or without a P1’ proline. A P1’ proline is required at the MA/CA cleavage site due to its role in a post-cleavage conformational change in the capsid protein. However, the HIV-1 PR prefers to have large hydrophobic amino acids flanking the scissile bond, suggesting PR recognizes two different classes of substrate sequences. We analyzed the cleavage rate of over 150 iterations of six different HIV-1 cleavage sites to explore rate determinants of cleavage. We found that cleavage rates are strongly influenced by the two amino acids flanking the amino acids at the scissile bond (**P2**-P1/P1’-**P2**’), with two complementary sets of rules. When P1’ is proline, the P2 side chain interacts with a polar region in the S2 subsite of the PR, while the P2’ amino acid interacts with a hydrophobic region of the S2’ subsite. When P1’ is not proline, the orientations of the P2 and P2’ side chains with respect to the scissile bond are reversed; P2 residues interact with a hydrophobic face of the S2 subsite while the P2’ amino acid usually engages hydrophilic amino acids in the S2’ subsite. These results reveal that the HIV-1 PR has evolved bi-functional S2 and S2’ subsites to accommodate the steric effects imposed by a P1’ proline on the orientation of P2 and P2’ substrate side chains. These results also suggest a new strategy for inhibitor design to engage the multiple specificities in these subsites.

## Introduction

The Human Immunodeficiency Virus type 1 (HIV-1) protease (PR), encoded in the viral *pro* gene, utilizes two aspartic acid side chains coordinating a water molecule to hydrolyze a peptide bond. Each subunit of the homodimeric PR contributes one aspartic acid to create this active site [1-3]. Surrounding these aspartates is a channel within the enzyme where a 7-to-8 amino acid stretch of protein interacts with the PR to determine its suitability for cleavage, with these interactions occurring through a series of subsites within the protease along this channel [4-6]. The PR itself is initially embedded in the viral Gag-Pro-Pol precursor [7]. Upon dimerization of monomeric subunits embedded within a pair of precursor polyproteins, the enzyme gains functionality and completes a three-or four-step intramolecular processing sequence to free its amino-terminal ends [8-10]. The resulting increase in structural stability enables the PR to intermolecularly cleave the primary HIV-1 structural polyprotein, Gag, as well as free the reverse transcriptase (RT) and integrase (IN) enzymes from the residual intermediates of Gag-Pro-Pol [11, 12]. Cleavage of the five processing sites within Gag releases small mature virion proteins, matrix (MA), capsid (CA), spacer peptide 1 (SP1), nucleocapsid (NC), spacer peptide 2 (SP2), and p6, and the cleavage proceeds in a specific order [12-16]. Protein processing results in the maturation of the genomic RNA dimer [17], assembly of the pre-reverse transcription complex, and envelopment of the ribonucleoprotein core within a capsid cone [18]. Incomplete processing results in a non-infectious virus particle [19-22].

Each of the ten cleavage sites recognized by the HIV-1 PR within Gag and Gag-Pro-Pol has a unique amino acid sequence (Table 1). There are a few common features, for example a β-branched amino acid cannot occupy the P1 position [16, 23], the amino acids directly flanking the scissile bond (P1/P1’) favor hydrophobic amino acids [24], and the sequences occupy a conserved shape or substrate envelope [25, 26], which is maintained even after substrate and enzyme co-evolution in response to PR inhibitors [27-30]. Nevertheless, no clear patterns exist to explain the varied rates at which they are cleaved [31]. Addressing that question is complicated by the fact that suboptimal cleavage site sequences could be used to help in the sequential cleavage pathway, or conformational and/or post-cleavage functional determinants could limit the use of optimal amino acids at the cleavage sites [32-34]. An unusual feature of the retroviral protease family is the ability to cleave sites with or without a proline in the P1’ position [16] i.e. just downstream of the scissile bond (MEROPS https://www.ebi.ac.uk/merops/cgi-bin/peptidase_specificity). The ability to cleave next to a proline is necessitated by the requirement for an N-terminal proline in the mature capsid protein (CA) that forms an intramolecular salt bridge to stabilize the post-cleavage CA structure and enable formation of the mature CA lattice [35-40]. Thus, this cleavage event is part of the regulation of virion assembly. This has previously led us to suggest that PR cleavage sites fall into two classes, those with a proline at P1’ and those with a hydrophobic amino acid at P1’ [24]. Left unclear is whether determinants of substrate specificity differ between the groups, and if so, how the homodimeric PR achieves multiple specificities.

**Table 1:** Amino acid sequences of HIV-1 cleavage sites

| Site | Abbreviation | P4 | P3 | P2 | P1 | P1' | P2' | P3' | P4' |
| --- | --- | --- | --- | --- | --- | --- | --- | --- | --- |
| **Matrix/Capsid** | **MA/CA** | **S** | **Q** | **N** | **Y** | **P** | **I** | **V** | **Q** |
| **Capsid/Spacer Peptide 1** | **CA/SP1** | **A** | **R** | **V** | **L** | **A** | **E** | **A** | **M** |
| **Spacer Peptide 1/Nucleocapsid** | **SP1/NC** | **A** | **T** | **I** | **M** | **M** | **Q** | **R** | **G** |
| Nucleocapsid/Spacer Peptide 2 | NC/SP2 | R | Q | A | N | F | L | G | K |
| **Spacer Peptide 2/p6** | **SP2/p6** | **P** | **G** | **N** | **F** | **L** | **Q** | **S** | **R** |
| Transframe (internal)/Transframe | TF*/TF | D | L | A | F | P | Q | G | K |
| **Transframe/Protease** | **TF/PR** | **S** | **F** | **S** | **F** | **P** | **Q** | **I** | **T** |
| Protease/Reverse Transcriptase | PR/RT | T | L | N | F | P | I | S | P |
| Reverse Transcriptase/RNase H | RT/RTH | A | E | T | F | Y | V | D | G |
| **RNase H/Integrase** | **RTH/IN** | **R** | **K** | **V** | **L** | **F** | **L** | **D** | **G** |

Here we report an extensive mutational analysis of six different HIV-1 PR cleavage sites using a previously developed two-substrate protease reaction to compare relative rates of cleavage [31]. All cleavage sites were held within the same context within a globular protein, and their initial reaction velocities were measured under near-physiological conditions. The measured initial reaction velocities were compared across the reactions through the use of an internal control protein substrate as a reference with a dynamic range of 1,000-fold for measuring rate differences. In total, 152 substrates were evaluated to define how sequence influences cleavage rate among the sites. We discovered that the PR binds the two classes of cleavage sites, as defined by the P1’ amino acid, in two different ways. The orientations of the P2 and the P2’ amino acid side chains within the S2 and S2’ subsites differ depending on whether or not a proline occupies P1’. This directional difference determines whether the subsites preferentially accommodate a hydrophobic or hydrophilic residue, an interpretation consistent with our previous structural analysis of substrates bound to a catalytically inactive PR [25, 26, 41]. The MA/CA and SP1/NC cleavage sites appear to be optimized for cleavage rate for the two classes of cleavage sites, while other sites represent suboptimal examples of these two classes of cleavage sites whose rates can be increased by manipulation of the P2 or P2’ amino acid. Thus, the HIV-1 PR has evolved bi-functional S2 and S2’ subsites to accommodate changes in side chain orientation in the P2 and P2’ substrate side chains imposed by the requirement to have a subset of cleavage sites with a P1’ proline.

## Results

### Two-substrate proteolysis allows accurate cross-reaction comparisons

For investigation of HIV-1 PR sequence specificity, we developed a system to compare the processing rate of cleavage sites under near-physiological conditions and independent of context. Efficient cleavage near neutral pH requires a globular substrate[31]. Six of the ten HIV-1 PR cleavage sites were investigated in this study. Briefly, the full MA/CA region of Gag served as an internal control in the two-substrate proteolysis system. A modified substrate (GMCΔ) was made from MA/CA by adding glutathione S-transferase (GST) at the N-terminus of MA and truncating C-terminal domain of CA to increase the size differences in the substrates and the cleaved products after proteolysis between the two substrates. A binding site for the Lumio Green Reagent (Invitrogen) was introduced into the Cyclophilin A loop of CA within both proteins, MA/CA and GMCΔ, to improve the detection of low-abundance cleavage products [42, 43]. Processing of the MA/CA cleavage site within GMCΔ was as efficient as the processing of the MA/CA cleavage site within MA/CA indicating that the alterations made to the GMCΔ do not affect the rate of cleavage of the MA/CA site (Supplementary Figure 1a). Five other HIV-1 cleavage sites (shown as bold in Table 1; CA/SP1, SP1/NC, SP2/p6, TF/PR [TF: Transframe], and RTH/IN [RTH: Reverse Transcriptase and RNase H, IN: Integrase]) were placed into the P4-P4’ positions of the cleavage site between the MA and CA domains of GMCΔ, each replacing the homologous MA/CA cleavage site. Although context was constant between all of the transplanted sites, we further minimized the influence of context by flanking the P4-P4’ residues with three glycines on each side (GMCΔgly). This caused a modest and uniform 3-fold decrease in the cleavage rate for all sites except for TF/PR which exhibited a 15-fold decline, suggesting a greater interaction with amino acids beyond P4 and/or P4’ for this site (Supplementary Figure 2). The control MA/CA substrate was likewise modified to include glycine spacer regions flanking the cleavage sites (MA/CA-gly). Each reaction was run with equal starting concentrations of GMCΔgly and MA/CA-gly so that the cleavage rate of GMCΔgly test substrates could be measured relative to the internal MA/CA-gly control substrate. We previously utilized this system to determine the relative processing rates of the wild-type cleavage sites from the Gag and Gag-Pro-Pol polyproteins [31].

### Only 11% of substrates in the library improved processing more than 3-fold

In total, 146 mutant HIV-1 cleavage sites were tested based on the six wild-type sites (Figure 1a, Supplementary Table 1). Most mutant sites differed from their wild-type parent by a single amino acid substitution, although 14 were dual substitutions; the substitutions chosen for each site were designed using the diversity of other sites as a guide. With the exception of the SP2/p6 and TF/PR cleavage sites, whose wild-type sequences were inefficiently cleaved, we also tested an alanine substitution in each position. Each site had every position mutated at least once, and 43 of 48 positions were tested with at least two different substitutions.

**Figure 1:**
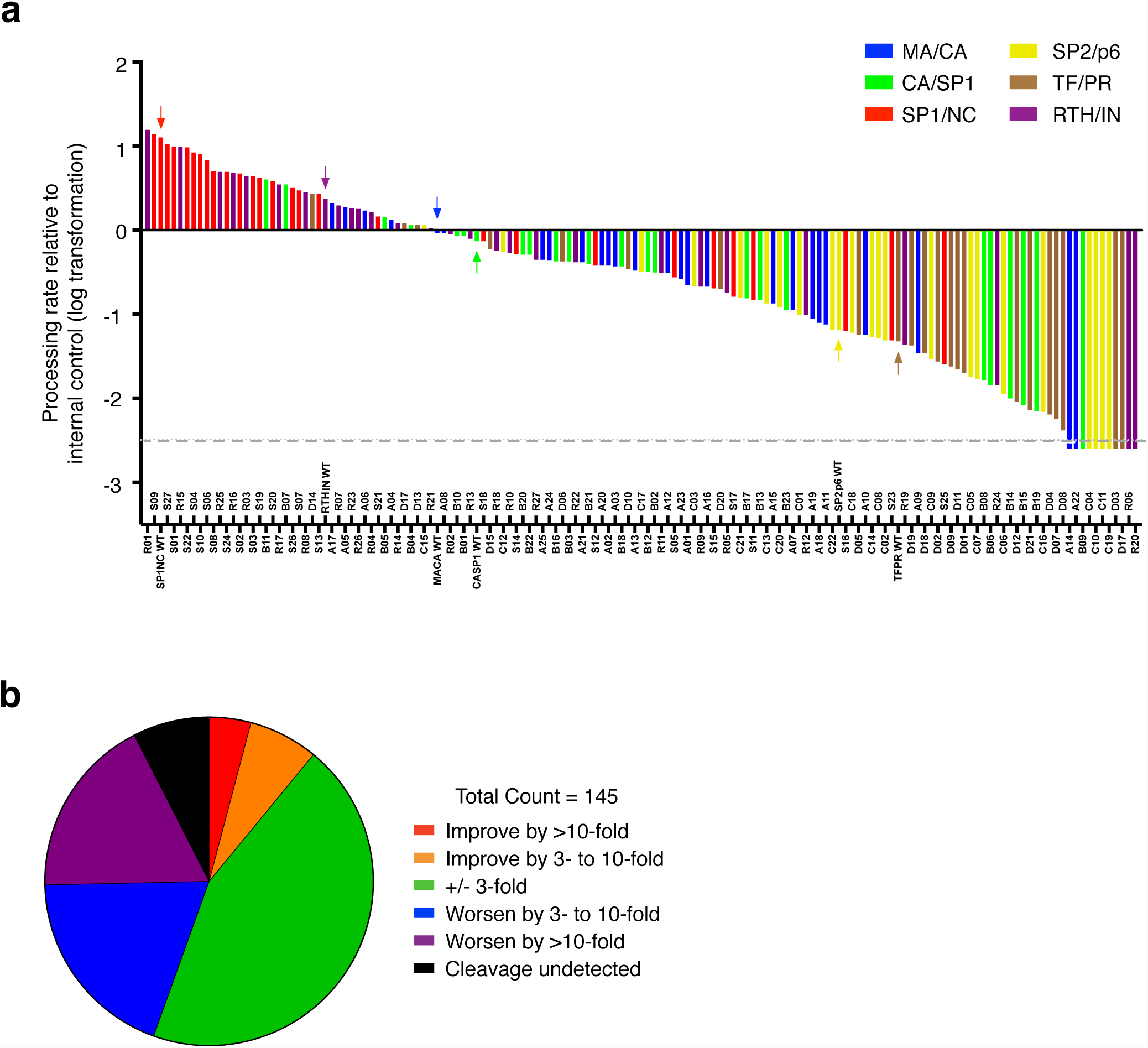
Relative processing efficiency of the altered substrates. (a) Distribution of GMCΔgly substrate cleavage efficiencies relative to the MA/CA-gly internal control substrate. The relative rates of substrate sites with wild type HIV-1 Gag sequences are indicated with arrows colored according to the figure insert. The dashed grey line represents the lower limit of detection in an extended 2h proteolysis assay. All bars extending below this line indicate substrates for which we did not observe cleavage. Site sequences and precise rate values are located in Supplementary Table 1. (b) Distribution of mutant substrate cleavage rates relative to their respective wild-type site.

The most common outcome of these mutated sites (65/146) was a cleavage rate change within 3-fold of their respective wild-type sequence (Figure 1b). We have largely limited the analysis of rate changes to those greater than 3-fold to focus on the larger effects even though under most circumstances the assay was accurate in measuring changes below this level of difference. Of the remaining 81 mutations, 28 slowed the rate of cleavage by 3-to 10-fold, and 26 by greater than 10-fold. Only 16 of the mutant sites tested improved the rate of cleavage over the parent site; those conferring a 10-fold or greater increase occurred only with substitutions within the inner four positions of the cleavage site (P2-P2’).

A small fraction of mutant substrates (11/146) had undetectable levels of cleavage, even when examined in an extended assay. Like the mutations that had the greatest positive impact on cleavage rate, these mutations were mostly within the central amino acid positions P2-P2’. Outside of this inner region, only mutations at the P3 position also ablated cleavage. However, these P3 cleavage-negative mutations were in the background of the SP2/p6 and TF/PR cleavage sites where the slow initial rate of processing for these two sites limited our ability to detect decreases in their efficiencies of cleavage beyond 20-fold. Further inspection of the patterns of changes in rates among the different cleavage sites ultimately revealed common features, as described in the next section.

### A working model for HIV-1 protease specificity

Interpretation of the results below will reference a working model that emerged while analyzing the data. The model suggests the existence of two prototypic sequence motifs: NΩ/PI (Asn, mainly Phe or Tyr /Pro, Ile) and βΦ/ΦE (β-branched aliphatic,hydrophobic/hydrophobic, Glu). When optimized for cleavage rate, both motifs form critical hydrogen bonds with residues 29 and 30 of *one* of the PR subunits. In the βΦ/ΦE motif, the P2’ glutamic acid (or glutamine) forms these hydrophilic interactions in the S2’ subsite; the P1’ proline in an NΩ/PI site shifts this interaction to the P2 asparagine in the S2 subsite. The reciprocal P2/P2’ residues are not involved in forming hydrogen bonds, but rather are β-branched amino acids, commonly isoleucine or valine, and they participate in distinct hydrophobic interactions with PR residues within the same S2/S2’ subsites. The data presented below show that the MA/CA and the SP1/NC sites are largely optimal for cleavage rate for the NΩ/PI and βΦ/ΦE sites, respectively, and that the remaining four sites can be manipulated to increase or decrease the rate of cleavage based on similarity to one or the other motif. One variation on this theme is that the size of the P1’ amino acid in the βΦ/ΦE influences the need for an optimal amino acid in P2’.

### MA/CA (SQNY/PIVQ): optimized for cleavage rate given a P1’ proline

MA/CA typifies the NΩ/PI motif due to its distinctive requirement for a P1’ proline in a post-cleavage conformation change. Proline as a constrained amino acid is a challenging substrate for most proteases such that no other proteases are known with mixed substrate specificity that includes a P1’ proline (MEROPS https://www.ebi.ac.uk/merops/cgi-bin/peptidase_specificity). Despite the presence of a P1’ proline in the MA/CA site, this site is cleaved relatively efficiently by the HIV-1 PR (Figure 1a), at about one-tenth the rate of cleavage of the most rapidly cleaved substrate. Furthermore, no single mutation within the key P2-P2’ inner region significantly improved the cleavage rate, not even substitution of the proline (Figure 2a). These data support the argument that the amino acid composition of the MA/CA site is essentially optimized given a P1’ proline.

**Figure 2:**
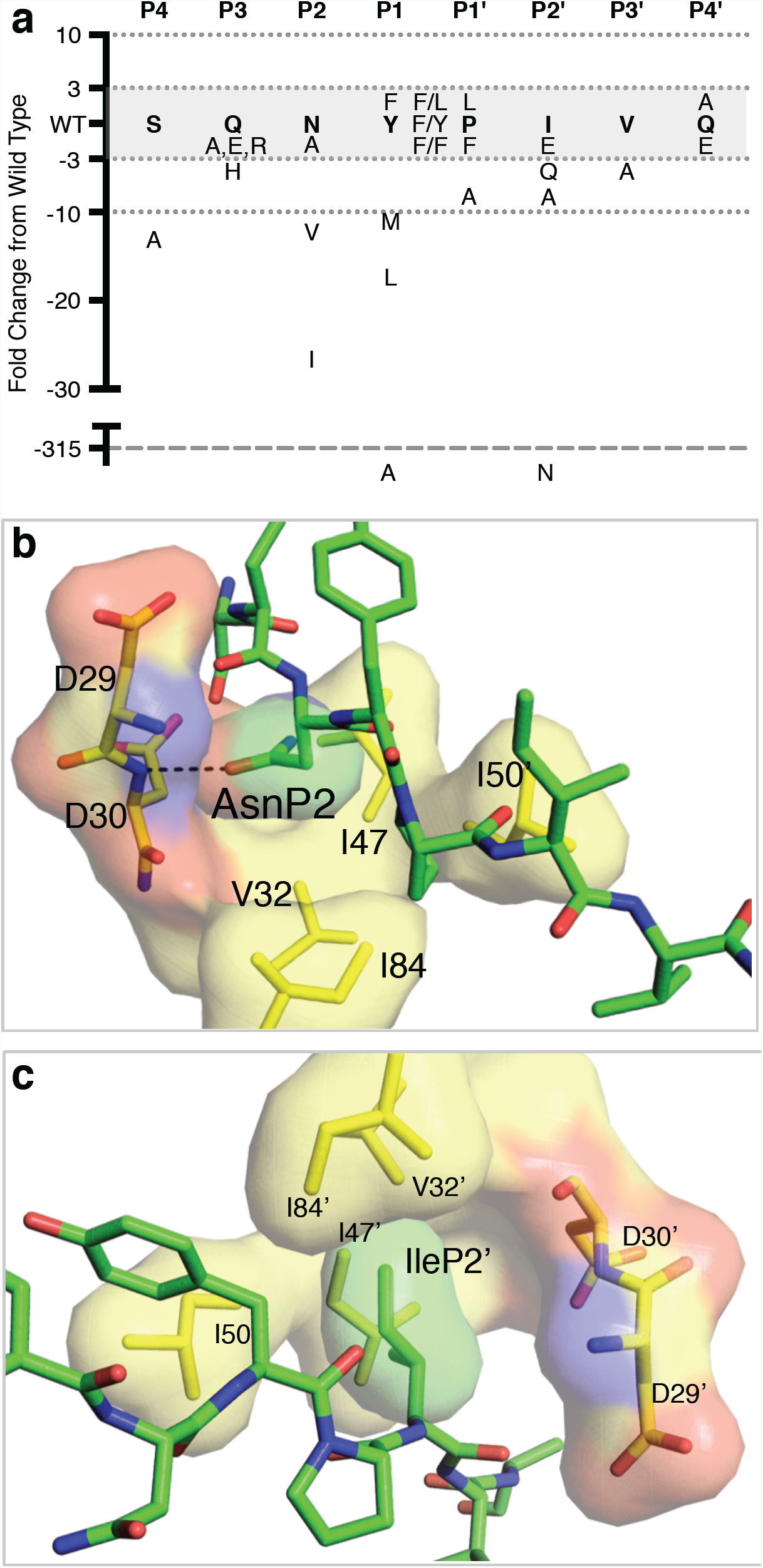
The MA/CA site represents the prototypical NΩ/PI site. (a) Relative change in processing efficiency for each mutant of the MA/CA site. Bold text denotes the wild-type residue at the indicated position. Amino acids within the greyed region exhibited +/-3-fold changes in cleavage rate. Amino acids located within the regions bordered by dotted lines exhibited between 3-and 10-fold changes in cleavage rate. Single amino acid substitutions with equivalent rates are separated by commas. Dual substitutions of adjacent residues are represented located halfway between columns and separated by a “/”. The dashed grey line represents the lower limit of detection for processing at this site. (b) Positioning of the P2 amino acid in the S2 subsite and (c) P2’ amino acid in the S2’ subsite. Amino acids from the HIV-1 PR are shown in yellow and labeled with their single-letter code and residue number in the PR. Primed versus unprimed residue numbers denote different PR subunits. Residues from the substrate are in green and labeled by three-letter code and positioning relative to the scissile bond. For both, nitrogen atoms are blue, oxygen atoms are red, and dashed lines denote hydrogen bonds. PDB file: 1KJ4.

While no P2-P2’ substitutions significantly enhanced the rate of MA/CA cleavage, several were deleterious. β-branched amino acids in P2 had strongly negative effects despite their frequent appearance at P2 in other sites, supporting a link between a P2 asparagine and the P1’ proline. Structural studies of a MA/CA peptide bound to PR (PDB: 1KJ4 [25]) showed the P2 asparagine side chain extends in the N-terminal direction (relative to the substrate) to make a hydrogen bond with the backbone of PR at the position connecting Asp29 and Asp30 (Figure 2b), an interaction that would not occur with a hydrophobic P2 amino acid. A P1 phenylalanine or tyrosine was more active than a P1 methionine or leucine (Figure 2a), as previously reported [16, 24, 44], suggesting optimal cleavage of P1’ proline sites requires a large aromatic amino acid in P1. In contrast to P2, an asparagine in P2’ was highly unfavorable (Figure 2a). The wild type amino acid in P2’, isoleucine, yielded the most rapidly cleaved site, though glutamine or glutamic acid were well tolerated. Figure 2c shows that the P2’ isoleucine angles away from Asp29’ and Asp30’ in the S2’ subsite, instead forming van der Waals (vdW) contacts with the hydrophobic amino acid Ile84’. Taken together the data point to a P1’ proline having an optimal cleavage rate with the motif NΩ/PI.

Most substitutions beyond P2-P2’ caused minimal changes in the rate of cleavage. However, one substitution, P4 serine to alanine, did have a significant negative effect, consistent with previous reports using peptide substrates [44-46]. This positive effect of this serine likely results from the ability of the P4 serine to form a hydrogen bond with the P2 asparagine when the side chain is oriented away from the scissile bond. Indeed, putting a P4 serine with P2 asparagine in a non-P1’ proline site had no effect on cleavage rate (see below, SP2/p6).

### SP1/NC (ATIM/MQRG): optimized for cleavage rate

Among the ten HIV-1 cleavage sites, SP1/NC is the most rapidly cleaved in its natural context [13-15], and could conceivably represent an optimal cleavage site sequence for the enzyme. Consistent with this possibility, our prior mutational analysis failed to significantly improve the rate of cleavage [16], and none of the 28 substitutions examined here did either. Among the other sites tested, only a mutated version of the RTH/IN site was cleaved faster than the wild-type SP1/NC sequence (Figure 1a), but at a rate that was only marginally better (1.2-fold).

Within the P2-P2’ core of SP1/NC (Figure 3a), substitution of P2 isoleucine with asparagine significantly reduced the rate of cleavage, demonstrating that the sequence requirements for the SP1/NC and NΩ/PI sites are distinct. This conclusion was reinforced by the β-branched valine substitution being well tolerated at the P2 position. Glutamine and glutamic acid gave equivalent rates as P2’ occupants, whereas small or β-branched alternatives resulted in varying degrees of reduced rates of cleavage, suggesting the non-NΩ/PI motif is based on a β_/_(E/Q) framework. Any of the common scissile bond residues (i.e. phenylalanine, methionine, and leucine) were active in either P1 or P1’, in agreement with previously published data [16]. The non-NΩ/PI motif therefore consists of βΦ/Φ(E/Q), in which Φ is a large, hydrophobic amino acid. The E/Q interchangeability was later resolved to favor glutamic acid, given it’s higher level of activity in the presence of a small P1’ residue (see the discussion of the CA/SP1 site below), yielding the βΦ/ΦE motif. Analysis of the structure of this substrate bound to the PR (PDB: 1KJ7 [25]) shows that the side chain of the P2 isoleucine in the βΦ/ΦE motif interacts with a hydrophobic pocket formed by Ile50 and Ile84 (Figure 3b). Meanwhile, the P2’ glutamine forms a hydrogen bond with the peptide backbone near Asp30’ of the PR (Figure 3b), and the amino-group side chain of the P2’ glutamine may additionally interact with the side chain of Asp30’ in this context; furthermore, the P2’ glutamine is oriented toward the C terminus relative to the substrate, pointing away from the scissile bond. In comparing the P2 and P2’ side chains in the two substrate-bound structures, their orientations are essentially reversed. This suggests the S2 and S2’ subsites have two recognition specificities, and are therefore bi-specific.

**Figure 3:**
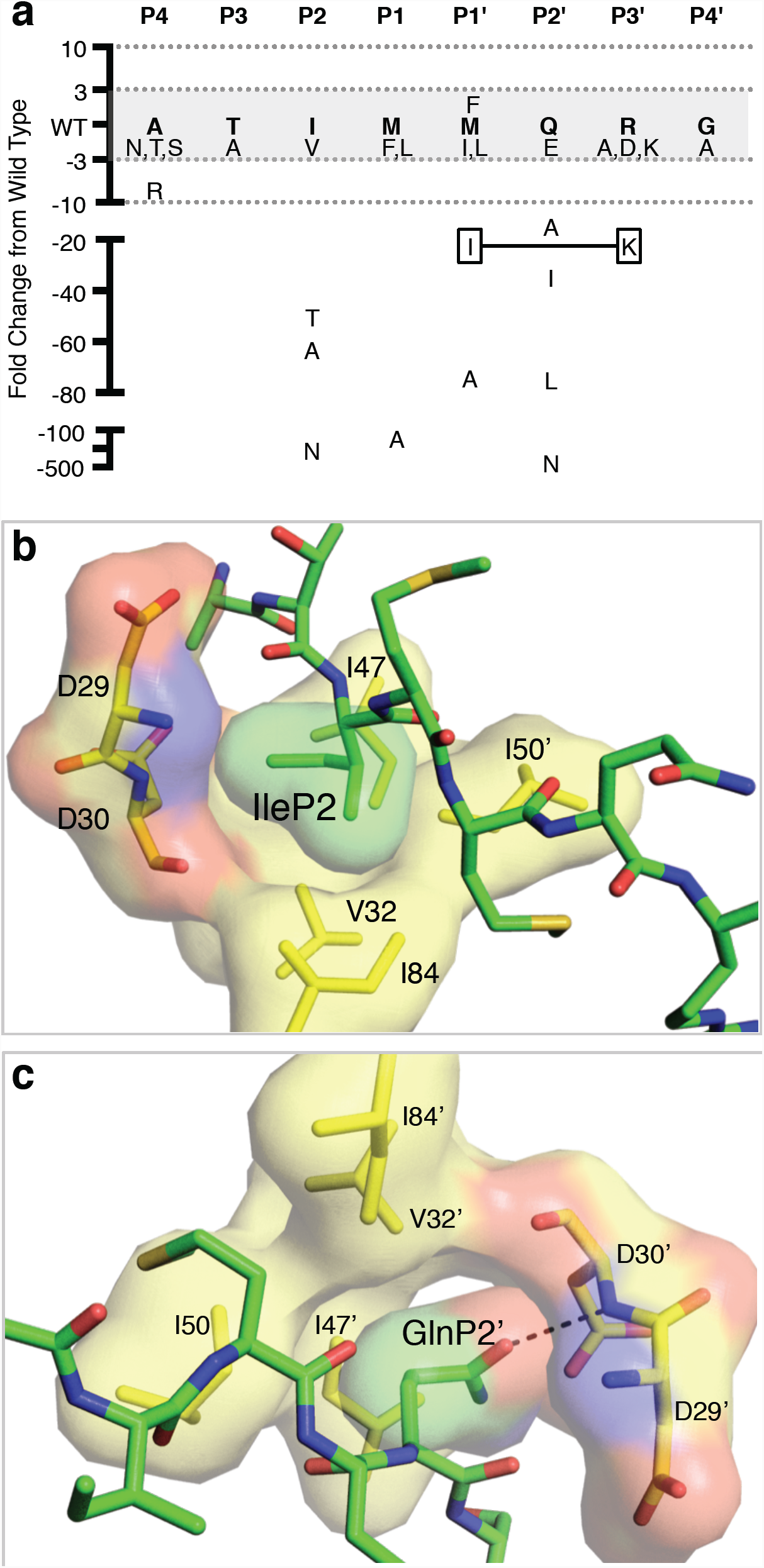
The SP1/NC site exemplifies an optimized βΦ/ΦE site. (a) Relative change in cleavage rate for each mutant of the SP1/NC site. All formatting is consistent with Figure 2, with the addition of a non-adjacent dual substitution mutant. These paired amino acids are boxed and connected by a solid line. (b) Positioning of the SP1/NC P2 amino acid in the S2 subsite and (c) P2’ amino acid in the S2’ subsite. Coloring is the same as in Figure 2b and 2c. PDB file: 1KJ7.

External to the P2-P2’ region, the most extreme effect observed was a 9-fold rate loss caused by an alanine to arginine substitution at P4. Although this again could suggest P4 as a potential rate-defining position, other P4 substitutions did not significantly affect the rate of cleavage of the SP1/NC site, implying arginine was the exception, not the rule. Moreover, arginine and alanine had only small effects in all other βΦ/ΦE sites.

### CA/SP1 (ARVL/AEAM): a βΦ/ΦE motif site with a small P1’ residue

The wild-type CA/SP1 sequence contains a β-branched P2 amino acid and a glutamic acid at P2’ identifying it as a βΦ/ΦE motif site, in addition to the absence of a P1’ proline. Accordingly, substitution of P2 to asparagine had a strongly deleterious effect (105-fold rate decrease), while isoleucine substituted for valine had a negligible effect. Moreover, all of the large common P1 amino acids were equivalently active substrates for the PR (Figure 4a). However, in contrast to SP1/NC where glutamic acid and glutamine were largely equivalent, the P2’ glutamic acid provided a substrate cleaved 60-fold faster than when P2’ was glutamine. The CA/SP1 core differs most significantly from the SP1/NC site at the P1’ position with alanine being the smallest hydrophobic/aliphatic amino acid. Examination of the entire dataset revealed a common preference in P2’ for glutamic acid over glutamine whenever P1’ was smaller than phenylalanine. This suggests that P2’ functions as a key compensatory position to improve PR recognition when the S1’ subsite is not fully occupied. An analysis of the crystal structure of this substrate bound to PR (PDB: 1F7A [25, 41]) (Figure 4b) shows that both the P2 and P2’ amino acid side chains are oriented similar to those of the SP1/NC site, consistent with the βΦ/ΦE motif.

**Figure 4:**
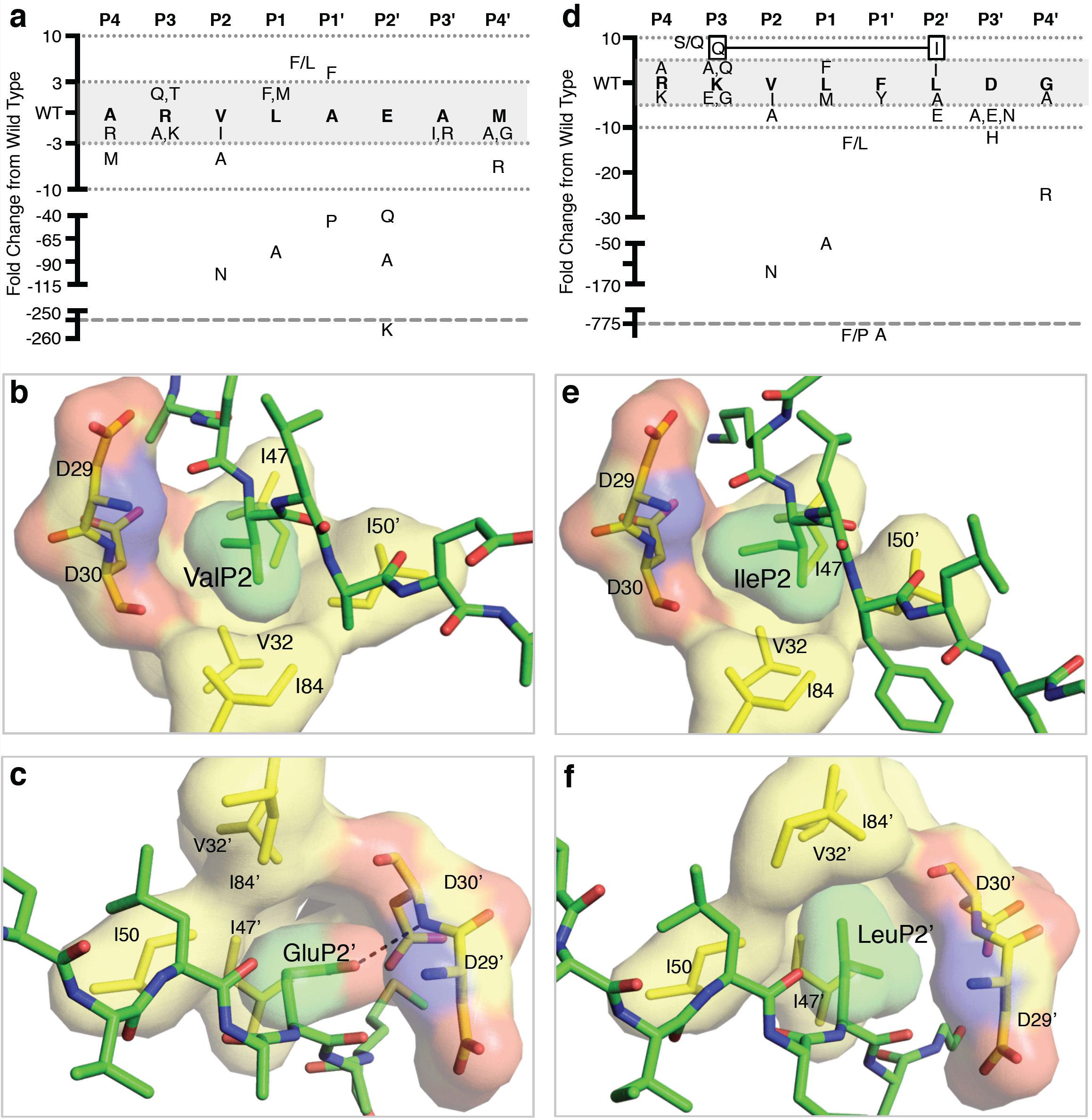
Suboptimal βΦ/ΦE sites demonstrate the compensatory role of P2’. (a) The relative change in cleavage rate, (b) positioning of the P2 amino acid, and (c) positioning of the P2’ amino acid of the wild-type CA/SP1 cleavage site. The RTH/IN site is represented in panels 4d, 4e, and 4f. Formatting and coloring of the figures remains the same as in Figure 2 and Figure 3. PDB files: (b,c) 1F7A; (e,f) 1KJH.

The external amino acids in CA/SP1 (i.e. P4, P3, P3’, and P4’), were all highly tolerant to substitution (Figure 4a). Among changes in size, hydrophobicity, or conformation of the R-group, only 2 of 11 substitutions changed the rate by greater than 3-fold and none exceeded a 7-fold change in rate. These results are consistent with previously published data [47], and support the idea that substitutions in these distal positions that elicit significant effects on processing rate are cleavage site-specific and not motif-dependent.

### RTH/IN (RKVL/FLDG): a βΦ/ΦE motif site with a large P1’ residue

With the absence of a P1’ proline and the β-branched valine in P2, the RTH/IN cleavage site belongs in the βΦ/ΦE motif. Mutagenesis confirmed this assignment; isoleucine and valine were equivalent in P2 while an asparagine substitution gave a poor substrate (Figure 4d). Furthermore, P1 accommodated large non-β-branched hydrophobic amino acids without affecting processing efficiency. An unusual feature in RTH/IN is the P2’ leucine; all P2’ aliphatic amino acids tested were moderately more active as substrates than glutamic acid. Comparison of the SP1/NC (Figure 3c) and RTH/IN structures (Figure 4f, PDB: 1KJH REF: [25]) shows similarity in positioning of both P2’ R-groups, although the terminal branched carbons of the leucine avoid positioning that would allow H-bond contacts with the Asp29’/Asp30’ backbone in favor of hydrophobic contacts seen in NΩ/PI sites. This ability to substitute these contacts was unique to the RTH/IN site, as a P2’ leucine in the SP1/NC context reduced cleavage efficiency nearly 80-fold. The large aromatic P1’ amino acid in the RTH/IN site could change the side chain angle of the P2’ amino acid to be more intermediate between the extremes of the two motifs allowing both conformations to be sampled for the P2’ side chain. Alternatively, an inverse relationship exists between the size of the P1’ amino acid and the importance of the P2’ position – the smaller the residue in P1’, the more important the identity of the P2’ position, and therefore which contacts are made in the S2’ subsite.

The non-core amino acids had measureable effects on the RTH/IN site. The P4 and P3 residues are both positively charged, and while individual substitutions had minimal effects on the rate of cleavage, co-substitution increased RTH/IN processing by 7-fold (Figure 4d). In the bound structure the P4 arginine occupies the S3 subsite and the P3 lysine occupies the S4 subsite [25]. This subsite swapping may be required to accommodate the size of the P4 arginine at the expense of reduced binding to PR. Conversely, five of the six substitutions of P3’ and P4’ tested slowed processing by at least 3-fold, with two of these substitutions having a greater than 10-fold effect. Taken together, the RTH/IN site appears subject to considerable site-specific effects on cleavage rate external to the core P2-P2’ region.

### SP2/p6 (PGNF/LQSR): a βΦ/ΦE motif site despite a hybrid amino acid sequence

The absence of a P1’ Pro suggests SP2/p6 should be a βΦ/ΦE motif site, but the asparagine at P2 is inconsistent with this designation. Isoleucine in P2 improved processing by 17-fold, while it had a minimally negative effect in P2’, behavior consistent with βΦ/ΦE motif sites (Figure 5a). That a P1’ proline measurably reduced processing of SP2/p6 compared to leucine or phenylalanine further supports the motif designation. Structural analysis of the side chain of the P2 asparagine in SP2/p6 shows its orientation mimics that of beta-branched amino acids in P2 when the P1’ amino acid is not proline (Figure 5b). This suggests that the nature of the P1’ amino acid determines the orientation of the P2 amino acid, in the case of SP2/p6 placing a polar asparagine into the hydrophobic face of the S2 subsite with the result being a suboptimal cleavage rate. Additionally, glutamic acid in P2’ enhanced processing, a finding consistent with an increased selectivity in P2’ when P1’ is non-aromatic.

**Figure 5:**
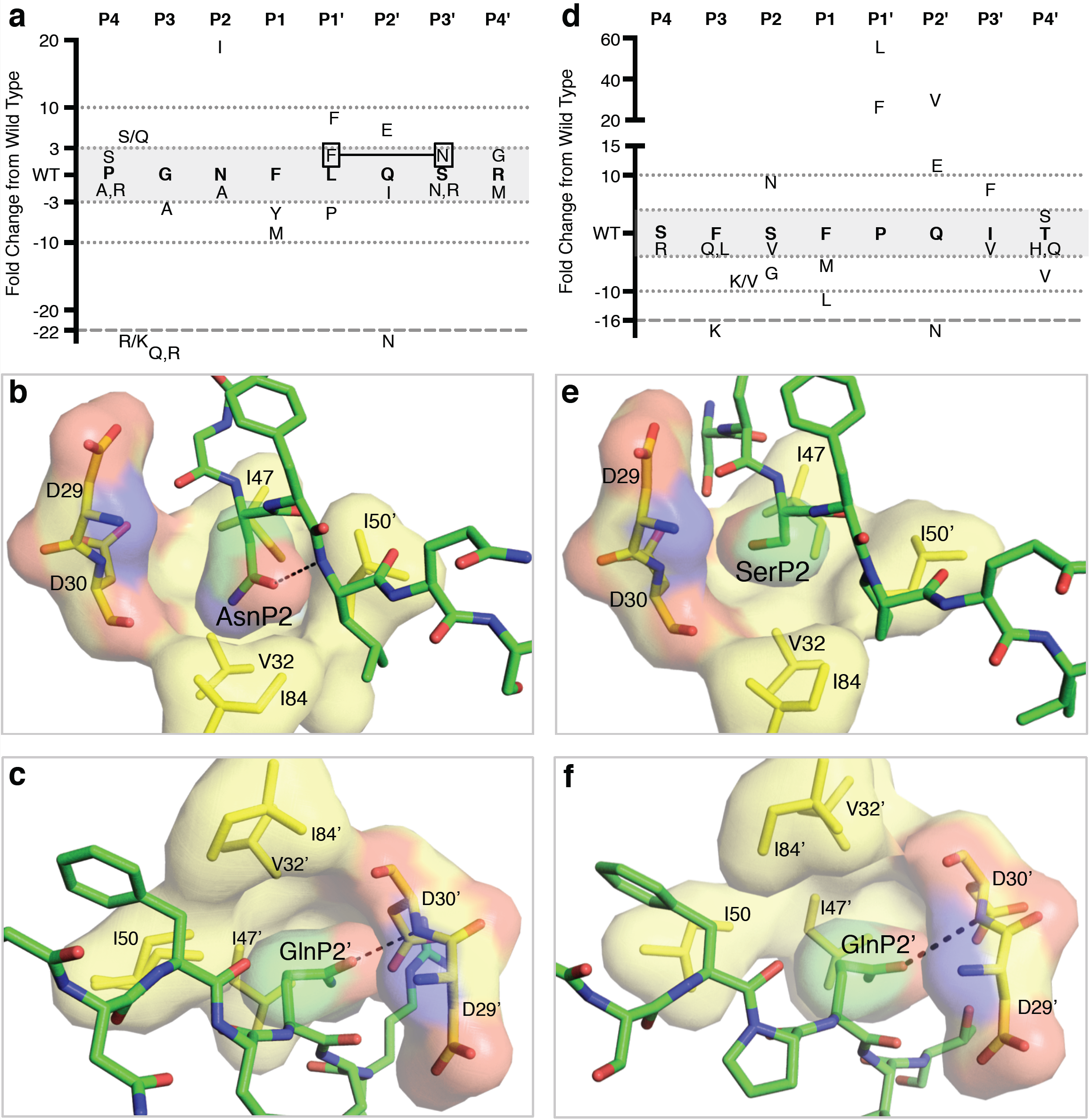
Hybrid cleavage sites have the lowest inherent rate of cleavage. (a) The relative change in cleavage rate, (b) positioning of the P2 amino acid, and (c) positioning of the P2’ amino acid of the wild-type SP2/p6 cleavage site. No structural data was available for the TF/PR site, so it was modeled in panels 4d, 4e, and 4f. Formatting and coloring of the figures remains the same as in Figure 2 and Figure 3. PDB files: (b,c) 1KJF; (e,f) 1KJ4 (enzyme only).

Among the distal sites, mutation of the P3 position with a large polar amino acid significantly affected SP2/p6 processing. Substituting arginine or glutamine into the P3 position resulted in SP2/p6 sites that were not detectably cleaved. The P4 proline creates a distinct conformation in which the P4 amino acid occupies the S3 subsite with the P3 glycine too small to fill the S4 subsite [25], and this is likely the reason for the added selectivity of P3. Consistent with this, replacing the P4 proline with serine rescues an otherwise deleterious P3 glutamine substitution (Figure 5a).

### TF/PR (SFSF/PQIT): a NΩ/PI motif site with several suboptimal amino acids

Like MA/CA, the TF/PR site contains a required P1’ proline, necessary in this case to restrict the set of cleavage sites available to the Gag-Pro-Pol–embedded PR [12] and placing this site TF/PR into the NΩ/PI motif grouping. In agreement with this designation, substituting an asparagine into P2 improved processing efficiency 9-fold, and a β-branched amino acid in P2’ conferred a 25-fold improvement over glutamine (Figure 5d). However, replacing the P1’ proline with phenylalanine or leucine improved the rate of cleavage by 24-and 57-fold, respectively. This is in contrast with the NΩ/PI site at MA/CA, in which substitution of the proline had minimal effect. This difference likely results from the absence of any other NΩ/PI-defining residues flanking the P1’ proline, effectively allowing the site to function as a βΦ/ΦE motif site when the proline is removed.

TF/PR resembled the majority of cleavage sites with regard to its flanking P4, P3, P3’, and P4’ amino acids. These outer positions contributed only modestly to determining the rate of cleavage. The sole outlier – P3 phenylalanine to lysine – resulted in undetectable levels of cleavage. This mutation could be rescued in part by a secondary substitution at P2, suggesting the observed effect was specific to the TF/PR site since arginine and lysine are often accommodated in this position. We additionally note that this, like the SP2/p6 P3 substitutions that blocked cleavage, occurred in the substrates with the slowest rate of cleavage. It may be that slow sites are proportionally more sensitive to mutations in their outer amino acid positions, or simply that we could not detect cleavage within the dynamic range of the assay.

Collectively these results reveal how the HIV-1 PR has evolved dual specificity to accommodate either the presence or absence of a P1’ proline. Such adaptation is necessary due to the fact that the proline alters the orientation of the P2 and P2’ side chains. By having S2 and S2’ subsites with dual recognition specificity the PR is able to define two distinct classes of substrate specificity.

## Discussion

We have used cleavage rates from over 150 distinct substrate sequences to explore the basis of HIV-1 PR cleavage site specificity. Since the initial separation of retroviral cleavage site sequences into those with and without a P1’ proline [24], there has been a question of whether substrate requirements would differ between the two families. Our data revealed the existence of two different substrate motifs, NΩ/PI and βΦ/ΦE, and that they are related to each other in a quasi-palindromic fashion in which important interactions with the N-and C-terminal halves of the substrate are switched between them. In βΦ/ΦE motif sites, a β-branched P2 residue occupies a majority of the S2 subsite, interacting with a region of hydrophobic amino acids while avoiding a more polar region outlined by PR residues Asp29 and Asp30. However, a P1’ proline alters the substrate interaction with the S2 subsite [48], reorienting the P2 side chain to face away from the scissile bond and into the hydrophilic region in the distal portion of the S2 subsite. There appears to be an opposite effect on the P2’ amino acid where the presence of a P1’ proline alters the positioning of the P2’ amino acid side chain that favors interaction with the equivalent hydrophobic region of the S2’ subsite, while the absence of a P1’ proline favors polar interactions with Asp29’ and Asp30’ in the distal portion of the S2’ subsite.

In generating our data, we engineered a system that used a globular protein substrate under near physiological conditions, and included an internal control substrate to act as a reference point from which we could reliably compare rates across reactions. A number of prior studies have examined HIV-1 PR substrate specificity [16, 24, 31, 34], though the vast majority of these made use of variably sized peptides under high salt and low pH conditions [44, 47-56], neither of which represent conditions likely encountered by the enzyme during virion maturation [57]. Moreover, these studies often mutagenized only a single target sequence (most often MA/CA or CA/SP1). Certain conclusions derived from these previous studies were found in our current study, namely the general flexibility in the external P4, P3, P3’, and P4’ positions and the critical role of P2 [46, 49-51]. Other conclusions appear to be incomplete due to the limited number of cleavage sites examined. For example, multiple studies reported glutamic acid to be the ideal P2’ residue [47, 48, 53]; in contrast, we found the specificity of the P2’ amino acid to be context dependent. Not only was glutamic acid suboptimal to β-branched amino acids in NΩ/PI motif sites, but its selectivity inversely correlated with the size of the P1’ residue in βΦ/ΦE motif sites.

The existence of two motifs raises the question of why HIV-1 doesn’t evolve to utilize one or the other. We aligned approximately 1100 HIV-1 subtype B Gag sequences to look for patterns of variability (Supplemental Figure 3) and found significant conservation at amino acid positions where a replacement could vastly improve the rate of cleavage in our assay (e.g. P2 asparagine in SP2/p6; Figure 5a). Considering the prolines of NΩ/PI sites have known functional roles outside of substrate recognition, this high level of conservation in βΦ/ΦE sites may similarly identify residues that take part in critical non-or post-cleavage functions. Indeed, CA/SP1, a βΦ/ΦE motif site, assumes an alpha-helical conformation during assembly to stabilize the immature capsid lattice [58-60], and mutagenesis of the site disrupted formation of virus-like particles [61]. Conversely, the fact that the observed variability that is present in cleavage site sequences would not change the order (meaning the intrinsic rate of cleavage) in which cleavage sites are cleaved implies selection for specific cleavage rates to temporally control certain processing events during virion maturation. While there is general agreement about the order of cleavage during maturation, the extent to which the kinetics of the cleavage events can be altered without affecting infectivity remains unexplored.

In this study we have focused on the primary sequence of the cleavage sites. While general features of the rate of cleavage are reproduced based on the primary sequence, it is clear that at least for some sites there will be additional rate determinants based on the context of an intact Gag or Gag-Pro-Pol precursor. For example, despite being a relatively poor substrate based on the cleavage site sequence alone, the SP2/p6 site is cleaved relatively efficiently in the context of a wild-type Gag molecule suggesting additional local determinants of cleavage [10, 31]. In some cases the P5’ position of the SP2/p6 site can be mutated in the context of selection for high level resistance to a PR inhibitor (along with the P1’ site) [62, 63], suggesting a role for context beyond the substrate binding cleft of the PR for at least some sites.

Finally, a review cleavage sites for other retroviruses can show if this pattern of use of the P2 and P2’ amino acids is conserved. The presence of a polar amino acid in P2 paired with an aliphatic amino acid in P2’ at the cleavage site that releases the N-terminal proline of CA appears to be relatively conserved among lentiviruses but not beyond [24]. The more generic pattern at the CA cleavage site for retroviruses is one where both the P2 and P2’ amino acids are hydrophobic, likely pointing to a more uniformly hydrophobic S2/S2’ subsite for these viral proteases.

Our work has provided fundamental insight into the nature of the protein code for HIV-1 PR cleavage sites as defined by the presence or absence of proline in P1’. The HIV-1 PR is an example of how the retroviral proteases are able to carry out the unique process of recognizing cleavage sites with or without the presence of a P1’ proline. This insight provides a strategy for systematically changing the rate of cleavage of sites, which will allow an analysis of the role of PR cleavage rate in these sites in the process of assembly and as a determinant of virion morphology and infectivity. In addition it is now possible to conceptualize inhibitor designs that could engage both specificities of the S2 and S2’ subsites.

## Materials and Methods

### Constructs

For the internal control substrate, the genomic sequence spanning the MA and CA domains was amplified from the pBARK plasmid, which contains the full *gag* and *pro* genes of the HIV-1 laboratory strain NL4-3, and subcloned into the pET-30b vector. The sequence was modified to include an N-terminal 6xHis tag, a tetracysteine motif (CCPGCC) in the Cyclophilin A binding loop (His87-Ala92) of CA, and three glycine residues between the P5-P4 and P4’-P5’ amino acids of the MA/CA cleavage site separating the domains.

The GMCΔ substrate was similarly derived from pBARK. However, in addition to the 6xHis tag, the tetracysteine motif, and the glycine residues, a GST domain was added between the 6xHis tag and N-terminal end of MA, and the CA region was terminated at position 278. These changes enabled detection by size of four different tetracysteine-containing proteins in a single reaction (the two substrates and their CA-containing products). All changes to the P4-P4’ region of GMCΔ were made via PCR-based site-directed mutagenesis. Primers were obtained through Sigma-Aldrich.

### Expression and purification of the HIV-1 PR and globular HIV-1 PR substrates

*Escherichia coli* BL21 DE3 lysogens (Novagen) were transformed with the various plasmid derivatives encoding the substrates, and grown in MagicMedia (Invitrogen) for protein production. Cells were pelleted and frozen at -80°C. After thawing, lysis was performed by sonication in TBS pH 7.5, 1% Triton X-100, 2 mM beta-mercaptoethanol. Cellular debris was collected by centrifugation, and the His-tagged proteins were purified by affinity chromatography using the Ni-NTA Superflow columns (Qiagen). Purified proteins were concentrated using Vivaspin Concentrators (GE Healthcare), with buffer exchange into storage buffer (20 mM sodium acetate, 140 mM sodium chloride, 2 mM beta-mercaptoethanol, 10% glycerol, pH 6.5). The pH was confirmed to within 0.2 units with a micro-pH electrode (Thermo Scientific).

Purified HIV-1 PR was produced as described previously [64, 65]. Briefly, HIV-1 PR was expressed from a pXC35 *E. coli* plasmid vector. The cell pellets were lysed and the PR was extracted from inclusion bodies with 100% glacial acetic acid. The PR was separated from higher molecular weight proteins by size-exclusion chromatography on a Sephadex G-75 column. The purified protein was refolded by rapid dilution into a 10-fold volume of 0.05 M sodium acetate buffer at pH 5.5, containing 10% glycerol, 5% ethylene glycol, and 5 mM dithiothreitol (refolding buffer).

### Two-substrate protease cleavage reactions

Two-substrate cleavage reactions were run in proteolysis buffer (50 mM sodium acetate, 50 mM NaMES, 100 mM Tris, 2 mM beta-mercaptoethanol, pH 6.5). Reactions were 150 μl in volume and pre-incubated at 30°C for 1 hour before addition of the enzyme to allow the Lumio Green Reagent (Invitrogen) to bind the CCPGCC motif in the CA region of each protein. Both substrates were included at an initial concentration of 1.2 μM. The HIV-1 PR was used at a concentration of 150 nM in the two-substrate assays where the substrates lack the glycine insertions, and 400 nM when the glycines were present to make up for the minor drop in processing rate. Aliquots were collected at specific time points throughout the course of the reaction and added directly to a tube with SDS to halt the reaction. The zero time point was removed immediately prior to the addition of enzyme. Most reactions were limited to 15 minutes, the timeframe required for the internal control to reach approximately 50% processing, though only data points generated within the first 10% of processing were used to generate relative rates. Sites exhibiting <10% cleavage after 15 minutes were retested in extended, 120-minute assays. If cleavage was still not observed after 120 minutes, the site was classified as not cleavable. After the final time point was collected, reaction pH was confirmed as 6.5 using a micro-pH electrode (Thermo Scientific). Substrates and products were separated by SDS-PAGE using precast 16% Tris-Glycine gels (Invitrogen). The fluorescently labeled proteins were then imaged with a Typhoon 9000 (GE Healthcare/Amersham Biosciences), and quantified by ImageQuant TL (GE Healthcare) software. The initial reaction rate for each substrate was determined using only the data points collected where the reaction was ≤10% complete, or was estimated based on the first non-zero data point collected. To determine the relative rate of processing, the ratio of initial velocities was compared using the internal control as the denominator, and the value recorded. The relative rate for each mutant was determined in at least two separate reactions. The average variance in estimated rate between the two reactions was 1.25-fold, with a range of 0 to 2-fold. Overall, these values differed by over 3000-fold, and were log-transformed for ease of interpretation.

## Acknowledgments

M.P. is supported, in part, by NIH Training Grant T32 AI07001. This work was supported by the NIH grant P01 GM109767. The work was also supported by the UNC Center For AIDS Research (NIH award P30 AI50410) and the UNC Lineberger Comprehensive Cancer Center (NIH award P30 CA16068).

